# Scaling use of the rust fungus *Puccinia punctiformis* for biological control of Canada thistle (*Cirsium arvense* (L.) Scop.): First report on a U.S. statewide effort

**DOI:** 10.1101/2023.09.30.560232

**Authors:** Dan W. Bean, Kristi Gladem, Karen Rosen, Alexander Blake, Robert E. Clark, Caitlin Henderson, John Kaltenbach, Joel Price, Emily L. Smallwood, Dana K. Berner, Stephen L. Young, Robert N. Schaeffer

**Affiliations:** Palisade Insectary, Colorado Dept. of Agriculture, 750 37 8/10 Rd, Palisade, CO 81526, USA; EcoData Technology, Plantsville, CT 06479, USA; Dept. of Biology, Utah State University, Logan, UT 84322, USA; Oregon Dept. of Agriculture, 635 Capitol St NE Salem, OR 97301, USA; USDA, ARS, FDWSRU, 1301 Ditto Avenue, Frederick, Detrick, MD 21702, USA; Retired, USDA, ARS, FDWSRU, 1301 Ditto Avenue, Frederick, Detrick, MD 21702, USA; USDA, ARS, Office of National Programs, Beltsville, MD 20705 USA

**Keywords:** Biological control, Canada thistle, *Cirsium arvense*, *Puccinia punctiformis*, systemic disease, thistle decline

## Abstract

Canada thistle (*Cirsium arvense* (L.) Scop., CT) is one of the worst weeds threatening temperate regions of the world. A host-specific rust fungus, *Puccinia punctiformis* (F. Strauss) Rohl., is known to cause systemic disease of CT, ultimately killing individuals and reducing stand densities. In 2013, it was demonstrated that fall inoculation of rosettes with coarsely ground leaves bearing *P. punctiformis* telia can successfully initiate epiphytotics. In the same year, a cooperative project between the Colorado Department of Agriculture and United States Department of Agriculture was initiated, in which CT patches across the state of Colorado (USA) were inoculated and tracked over subsequent years for changes in stem density. Here, we report our findings from 8 years (2014-2021) of monitoring effort. At most sites (*N* = 87), CT stem densities declined, from a mean (± SE) of 87.9 (± 6.5) stems to 44.7 (± 4.2). These declines however were spatially-autocorrelated, and likely attributable to local growing conditions, as mean annual daily maximum temperature and standard deviation of elevation, as well as climatic conditions around the times of both treatment and monitoring, were found to be important predictors of CT decline. Further, we observed that the amount of inoculum deployed, timing since last release, and method in which it was spread locally at a site were also associated with the magnitude of CT stem decline. These results are indicative of the value of *P. punctiformis* as a CT biological control agent. The name *Cirsium arvense* dieback (CADB) is proposed herein to describe the agriculturally important decline in CT stem densities attributable to this previously un-named systemic disease.

## 1. Introduction

Canada thistle, also known as Californian thistle and creeping thistle, (*Cirsium arvense* (L.) Scop., Asteraceae, CT) is a noxious perennial weed of temperate pastures, rangelands and agricultural lands worldwide (Moore, 1975; Morishita, 1999; Guiggisberg et al., 2012). Native to southeastern Europe and North Africa, CT is believed to have been introduced to North America from Europe in the 1600’s as a contaminant in seed grain (Guiggisberg et al., 2012). Canada thistle has since spread to 45 of the 50 U.S. states and 12 of the 13 Canadian provinces and territories. Recognized as a noxious weed as early as 1795 (Morishita, 1999), CT currently holds this designation in 43 states and 6 provinces, respectively (Kartesz, 2015). Where it has invaded, it causes economically measurable yield losses, often mediated through allelopathy in agricultural systems (Stachon and Zimdahl, 1980; Donald, 1994), as well as negative impacts on natural ecosystems (Cheater, 1992).

Canada thistle reproduces sexually by seed and vegetatively by shoot buds (Donald, 1994). Although seeds (achenes) are not as important as asexual reproduction in patch expansion, they are important for establishment of new patches (Donald, 1994). Canada thistle is imperfectly dioecious or subdioecious, with male and female plants tending to grow in patches consisting of a single sex, since patches often consist of one individual that has spread through extensive clonal growth (Kay, 1985). Cross pollination between male and female patches is necessary for seed production, although seed viability is generally low, and the initial seedling shoot does not produce flowers (Amor and Harris, 1975). Once a patch is established, vegetative spread through horizontal growth of the root system is rapid, especially when competition from other plants is low. As a result, CT typically grows in long-lived clonal patches from several meters up to 25 m or more in diameter (Donald, 1994). Canada thistle does not have rhizomes; rather it spreads vegetatively via thickened roots with adventitious shoot buds that grow both vertically and horizontally (Donald, 1994). Three types of underground structures are produced by CT: long, thick horizontal and vertical roots, short fine roots, and vertical underground stems (Hamdoun, 1970). All of these structures can produce buds which lead to new shoots.

The extensive root system found in CT patches makes the plant difficult to control. Traditional land preparation (plowing or tilling) aggravates CT infestations by cutting up roots and producing fragments which bud and produce new shoots and patches (Magnusson, et al., 1987). Herbicides that kill the root system are the primary option for CT control in row-crop agriculture (Beck, 2008), although this option is only moderately effective since CT plants frequently recover from treatment (Carlson and Donald, 1988; Thomas et al., 1994). Current recommendations for CT control involve repeated herbicide applications and mowing over several seasons (Beck and Sebastian, 2000; Beck, 2008), but these control methods are generally not economical, especially on lands of marginal value (Amor and Harris, 1977; Beck and Sebastian, 2000).

Effective biological control agents for CT are much sought-after. At least eight organisms, including fungi, bacteria, and insects have been evaluated for use as biological control agents of CT (Cripps et al., 2011, 2012), but these have had minimal impact, and are not regarded as effective (Price, 2014). Currently, the most promising biological control agent for CT is the obligate rust fungus *Puccinia punctiformis* (F. Strauss) Rohl. (hereafter *Puccinia; Puccinia sauveolens* (Pers.) Rostr., = *Puccinia obtegens* (Link) Tul.), which has long been naturalized in North America. The fungus is found in all states and provinces in which CT is found, and in 1893 became perhaps the first plant pathogen proposed as a weed biocontrol agent (Halsted, 1894; Wilson, 1969).

*Puccinia punctiformis* has a complex life cycle, highly adapted to its host (Cockayne, 1915; Buller, 1950; Menzies, 1953; Berner et al. 2013b). The fungus is an obligate biotrophic autoecious rust (completes its life cycle only on living CT plants) that can systemically infect individuals (French and Lightfield, 1990), likely resulting in permanent infection of the root system (Cockayne, 1915; Menzies, 1953). Some of the newly emerged shoots from systemically diseased roots are tall, spindly, deformed, and rarely flower with leaves bearing orange-colored haploid spermagonia (Fig. 1 A). These produce a nectar-like solution (Fig. 1 A), which contains haploid spores (spermatia) and has a pleasant fragrance that attracts insects (Connick and French, 1991; Naef et al., 2002). In turn, insects may carry the solution, bearing spermatia, from one spermagonium to another of a different mating type, on a different genet, resulting in outcrossing (Menzies, 1953). Following fertilization (fusion of newly transferred spermatia and receptive hyphae within the compatible spermagonium), spermagonia give rise to brown aecia which bear dikaryotic aeciospores (Fig. 1 A, B). The aeciospores spread to leaves of nearby CT shoots and produce uredinia and telia in late summer (Fig 1C, Berner et al., 2015b). Under favorable environmental conditions, the two-celled teliospores (Fig 1D), produced in the telia, germinate, undergo meiosis, and produce a basidium with haploid basidiospores (Berner et al., 2013b)). In addition to favorable environmental conditions, germination of the teliospores also requires factors produced by Canada thistle tissue (French 1990; Clark et al 2020) which ensures that germination and basidiospore formation most often occur on or in close proximity to the host plant. Teliospores that settle on newly emerged fall rosettes may germinate, and the resulting basidiospores are well positioned to infect the rosettes (Berner et al., 2013b), resulting in the growth of mycelia that extend into the roots and shoot buds on the roots (Menzies, 1953). Once roots are systemically infected, the fungus may remain belowground as mycelia within the root system for one to several years without initiating the conspicuous sexual portion of the life cycle (Buller, 1950; Menzies, 1953; Berner et al., 2015a). The visible reproductive portion of the fungal life cycle occurs above ground and includes showy and fragrant spore-bearing systemically diseased shoots, yet *Puccinia* exists primarily as a cryptic obligate root parasitic fungus (Olive, 1913) often making it difficult to track in the field.

**Figure 1.**
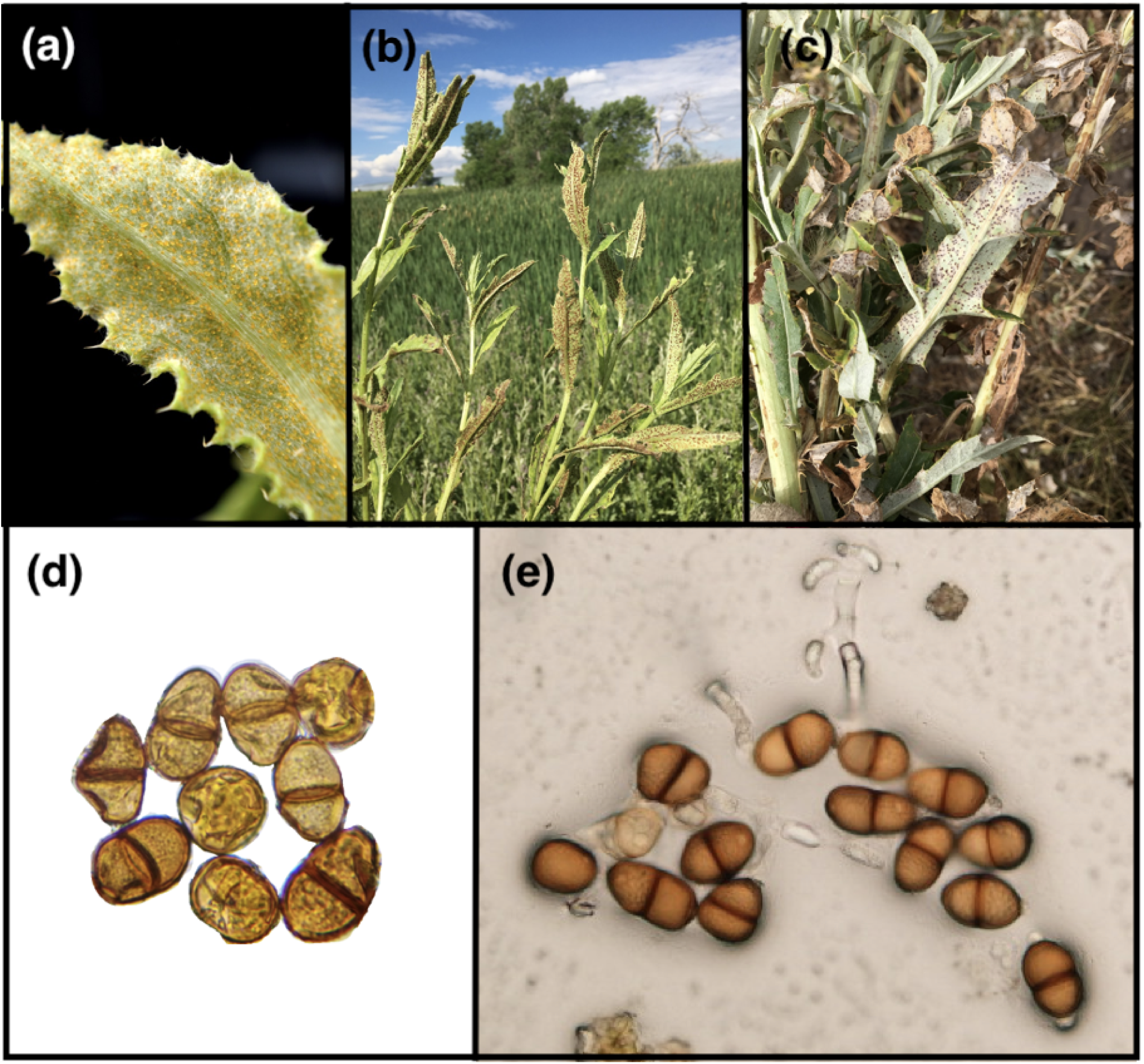
Characteristics of the Canada thistle – *Puccinia punctiformis* pathosystem. (A) Systemically infected leaf showing spermagonia, with orange nectar. (B) Systemically infected shoot bearing aecia with aeciospores. (C) Leaves of Canada thistle bearing telia of *Puccinia punctiformis* resulting from infection by aeciospores. (D) Two-celled teliospores, single celled uredinospores. (E) Two-celled diploid teliospores with a basidium and haploid basidiospores. Adapted from Berner et al. 2013b.

**Figure 2.**
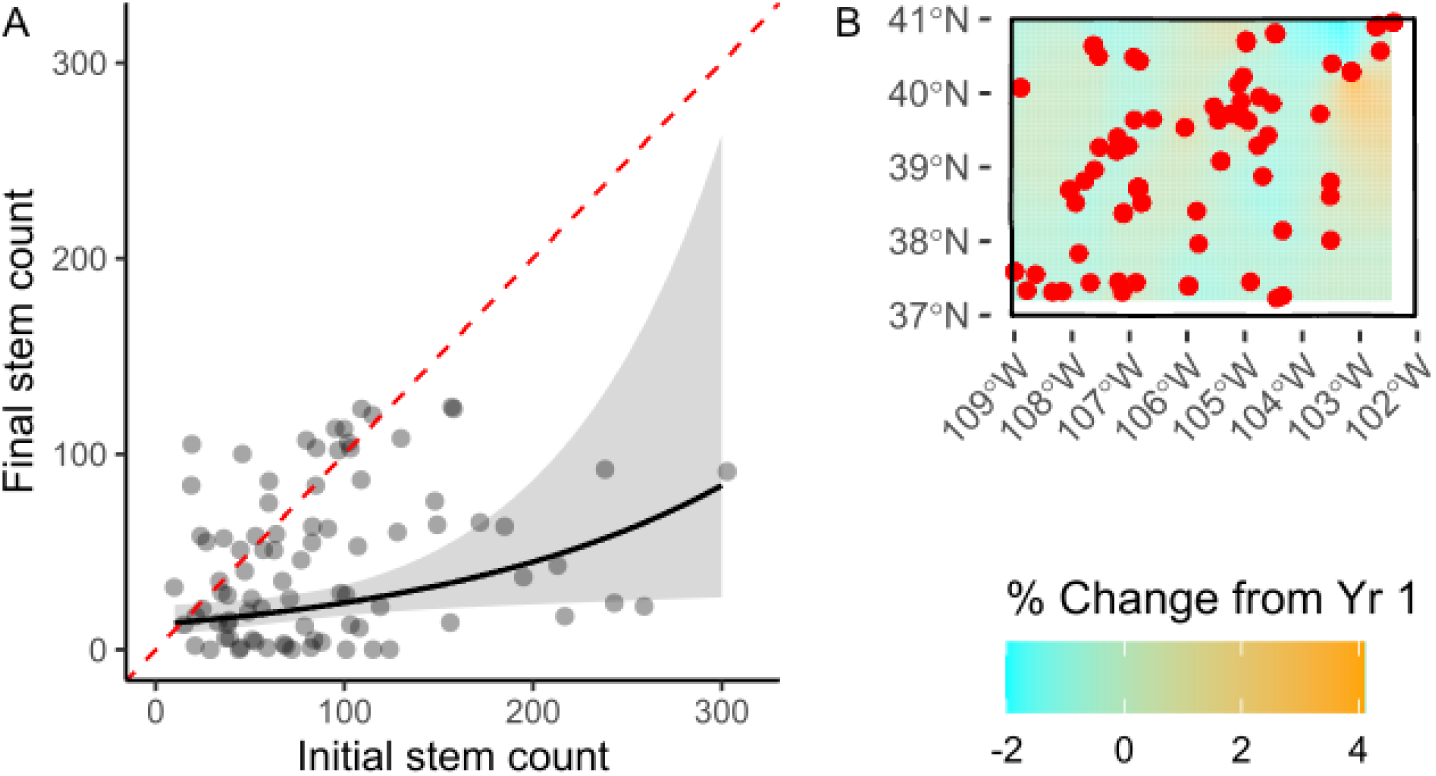
Initial vs. final Canada thistle stem counts. Points indicate sites (*N =* 87). Dashed red line represent y=x, with values above the line indicating an overall increase in stem count, and values below a decline. Shaded area is 95% confidence ribbon. Percent change (B) in Canada thistle stem counts from initial to final survey, interpolated across Colorado with ordinary kriging. Red points represent site locations across the state where thistle stands were treated with *Puccinia punctiformis* and subsequently monitored.

The potential for CT control using the rust has long been recognized, starting in 1893 (Halsted, 1894; Wilson, 1969) and continuing with Cockayne in New Zealand who proposed that while rust could be used for biological control, it was exceedingly difficult to collect enough teliospores for inoculation (Cockayne, 1915). Additional studies over the last century further revealed the potential for *Puccinia* to serve as a biocontrol agent. Ferdinandsen (1923) documented an increase in systemically diseased shoots over a three-year period in undisturbed pastures, reaching a maximum of 70% diseased shoots at the end of his study. Watson and Keogh (1980) documented the effects of *Puccinia* on mortality of the thistle and provided data on the destructiveness of rust infections. Thomas et al. (1994) showed that infection with *Puccinia* reduced both flowering and vegetative reproduction. Although different ecotypes of CT have been described (Moore, 1975), all ecotypes of CT tested by Turner et al. (1981) were susceptible to the rust. In sum, the documented effectiveness of natural infections in controlling CT has been both intriguing and frustrating, since establishment of rust fungus epiphytotics (outbreaks of a plant disease) has not been readily achieved by weed control practitioners. The difficulty in purposefully establishing epiphytotics has remained the primary barrier to development of an effective CT biological control method (Berner et al., 2013b).

Building on previous work, Berner et al. (2013b) refined understanding of the rust disease cycle and used the information to develop protocols to purposefully establish epiphytotics of the rust on CT patches. In these protocols, infective teliospores from telia produced in mid-to late-summer are deployed as inoculum, and inoculations are made utilizing fall-emergent rosettes as the infection court (plant stage most susceptible to systemic infection). With these protocols, rust epiphytotics have been successfully established by researchers in New Zealand, Russia, Greece and the U.S. After an initial round of inoculations at 10 sites, reductions in CT shoot densities averaged 43.1 ± 10.0% at 18 months after inoculation, 63.8 ± 8.0% at 30 months after inoculation, 80.9 ± 16.5% at 42 months after inoculation, and 72.9 ± 27.2% at 54 months after inoculation (Berner et al., 2013b). This work has revealed an inexorable, linear decline in thistle densities, requiring no other intervention than inoculation of fall rosettes with teliospores during study set-up.

The Colorado (CO) Department of Agriculture (CDA) has previously responded to heavy demand for CT biocontrol by redistributing the Canada thistle gall fly, *Urophora cardui* (L.) within state, through Colorado’s Biological Control Program (BCP). Since the fly had become established throughout CO, and impact on the target appeared to be insignificant, as noted by other biocontrol practitioners (Cripps et al., 2011; Price, 2014), the program was discontinued even though demand for CT biocontrol remained high. In 2013, a multi-year cooperative project between the CDA and United States Department of Agriculture, Agriculture Research Service (USDA-ARS) began to test the feasibility of using *Puccinia* in a statewide CT biological control program (Berner and Bean, 2013a). The objectives of the project were to: 1) Routinely establish epiphytotics of systemic disease, caused by *Puccinia*, in CT patches by inoculating fall-emergent rosettes with ground, telia-bearing leaves; 2) Improve efficiencies of inoculation methods (e.g., inoculation timing, amount of inoculum, etc.); 3) Train stakeholders in CO and surrounding states in establishing successful epiphytotics; 4) Monitor CT densities to document the impact of this biological control approach. This manuscript reports the results of these efforts.

## 2. Materials and Methods

### 2.1. Site selection, inoculation, and monitoring

Beginning in 2013, and continuing through 2021, a total of 181 CT-infested sites in CO were scouted for rust inoculation. The sites were primarily selected from a list of CO residents who had requested the Canada thistle gall fly, *U. cardui*, through the CDA’s BCP centered at the Palisade Insectary, Palisade, CO. The residents on the list were asked if they would like to participate in a multi-year monitoring study that used the rust fungus to control CT on their property. Other sites were selected in cooperation with county weed managers who were given information about *Puccinia* and the statewide trial program. Additional sites were volunteered by landowners who had attended workshops, talks and tours sponsored by the BCP, while outreach to organically certified producers resulted in the acquisition of several more study sites. Finally, an effort was made to establish sites in each of Colorado’s 64 counties in order to evaluate rust efficacy across a broad landscape and associated variation in both climatic and terrain conditions.

### 2.2. Inoculum collection and preparation

Telia-bearing CT leaves (Fig. 1 C) were used to inoculate all of the sites. The leaves were collected in September each year across CO. During the course of this study, numerous CT patches naturally infected with *Puccinia* were discovered. These were identified as potential teliospore collection sites. In mid-to-late summer of each year, these potential sites were surveyed for telia, which appear as small pustules on infected yellowing CT leaves (Fig. 1C). Leaves were harvested and stored in paper bags to allow foliage to dry at room temperature. Dried leaves were ground to a coarse powder in a kitchen blender and used as inoculum that season, or stored at –80°C for future use. Samples of ground leaf preparations were viewed under a microscope to ensure the majority of spores were two-celled teliospores (Fig. 1 D), the spore type necessary for initiating systemic infection (French and Lightfield, 1990; Van den Ende et al., 1987).

### 2.3. Inoculation timing and procedure

Inoculation timing was planned to coincide with optimal temperature and dew point for systemic infection when leaf-surface moisture was likely (Berner et al., 2013b). Averages of historical weather data were generated for daily dew points and maximum, minimum, and average daily temperature. Data, for as many years as available, were obtained online from weather stations closest to each site, and averages over years were calculated for each day of the year. Average dew point temperature and minimum air temperature were used to predict likelihood of dew for each day. Likelihood of dew was calculated as average dew point temperature minus minimum air temperature, with values greater than zero indicating likelihood of dew (Berner et al., 2013b). This likelihood, combined with average air temperature between 13 and 18° C, the optimum temperatures for teliospore germination, were the predictors of optimum inoculation day for each site. These calculations formed the target range of days for inoculation at each site, but inoculation dates did not always coincide with predicted times. Weather averages across CO are highly variable, in part due to differences in elevation (1000 to 4400 m above sea level). In some instances, there were no stations at the same elevation as inoculation sites. At such sites, inoculations always took place in fall after observed dew and new rosettes. Sites were inoculated at various times during the day, depending upon travel schedules and the availability of landowners. In each site, a 12 x 4 m transect was permanently marked using metal posts (see section 2.4 below for more detail). For sites with ∼40-50 rosettes along the transect, CT was inoculated by placing ∼0.5 g of inoculum in the crown of each rosette (Targeted). Handheld spray bottles were then used to apply a fine mist of water to the point of run-off, for the purpose of adhering the inoculum and creating a moist microclimate for spore germination. For sites with no (Broadcast) or limited rosette availability (Targeted +Broadcast), spores were hand distributed across plots, with the amount applied recorded. For sites with intermediate rosette availability, we used a combined approach. Finally, unless sites were damaged by livestock, fire, flood, herbicide application, or mowing, each patch in each site was inoculated once. We also inoculated select sites again if we did not find any evidence of systemic infection from the previous year’s inoculation.

### 2.4. Field plot establishment and monitoring protocol

Potential monitoring sites were evaluated to determine if aboveground signs of *Puccinia* were present prior to inoculation. Each site was visually assessed by walking through the CT infestation in search of leaves bearing spores, or systemically infected stems. If the rust fungus was not observed, we proceeded with the monitoring protocol. Our permanent monitoring protocol required a CT infestation a minimum of 12 m x 4 m and placement of two metal posts, positioned 12 m apart to mark the start and end of a transect with one post designated as the origin. Transect placement targeted healthy, dense stands near the center of a patch, and transect direction was determined with a handheld compass. A GPS point was recorded, and a photo-point was established from the origin post. Sites were monitored annually, between June and August, for evidence of systemic disease (i.e., spore-bearing systemically diseased shoots) and CT shoot density.

At each transect, a tape was extended 12 m from the point of origin post to the second post. A photo was taken sighting down the transect from the origin post. Monitoring began at two meters. A plot monitoring square, measuring 60 cm x 60 cm was placed to the right of the tape with the two-meter mark in the upper corner ensuring the same ground would be counted annually. Healthy and infected stems rooted in the square were each recorded. Monitoring continued along the meter tape every two meters until six plots were recorded. Weed stage and disease progression stage were recorded along with land-use.

### 2.5. Statistical analyses overview

The effects of biocontrol treatments, time, geography, and climate on CT stem counts were evaluated for 87 of the original 181 sites. Though a considerable reduction, these sites span 42 of Colorado’s 64 counties, and have at minimum three years of monitoring data. Reasons for site exclusion from the overall analysis include: less than three years of monitoring data, evidence of landowner intervention (i.e., herbicide application), or signs of systemic infection were already present at the site in year one. Effects were evaluated through Poisson generalized linear mixed effects models (GLMMs) and ordinary kriging. Due to data sparsity, only main effects were considered in the GLMMs, and dimension reduction was performed on multiple climatic variables using principal component analyses (PCA). All mixed effects models included site ID as a random intercept term.

*Overall analyses –* Overall trends from the beginning to the end of the time series were evaluated by modeling final stem counts as a function of initial stem counts, by site. Ordinary kriging was also performed on the percent change in stem counts from the first to the final survey to test for spatial autocorrelation. These initial results were used to inform the next analyses. In particular, ordinary kriging suggested that terrain ‘flatness’ may have influenced stem counts over the time series.

*Per-timestep analyses* – The change in stem counts from time *t*-1 to time *t* was also evaluated to account for inconsistent treatment protocols at each site. To do so, we used two different Poisson GLMMs (see: *Protocol effects* and *Site effects*). In both models we included stem count at time *t*-1 and survey year as covariates, to account for temporal trends and autocorrelation.

*Protocol effects* – The first model focused on treatment protocol, and included as predictors: treatment type at last treatment (Targeted, Broadcast, or Targeted+Broadcast), years since the last treatment, and amount of inoculum used in the last treatment.

*Site effects* – The second model focused on (site-level) climate and geographic effects, and included as independent variables: annual mean maximum temperature, standard deviation of elevation (within a 2.5 km radius around each site), % cropland or developed land use (within a 1 km radius around each site), and the first and second principal components extracted from two different PCAs.

*Principle component analyses* – The first PCA (henceforth ‘treatment-time PCA’) included climate variables from a 2-week window prior to each biocontrol treatment, while the second PCA (henceforth ‘observation-time PCA’) included the same variables but collected from a 2-week window prior to each monitoring survey. Both PCAs included the following predictors (based on daily means averaged over the 2-week window): maximum temperature (C), minimum temperature (C), average temperature (C), total precipitation (mm/day), water vapor pressure (Pa), relative humidity (%), dew point (C), shortwave radiation flux (W/m^2^), as well as elevation (m).

Briefly, in the treatment-time PCA, PC1 explained 52.1% of the variance while PC2 explained 22.8% (Fig. S3A). Higher values of PC1 correlated with higher elevation and precipitation, while lower values of PC1 correlated with higher temperature, vapor pressure, and dew point. Higher values of PC2 correlated with higher relative humidity and also correlated with precipitation, while lower values correlated with higher radiance.

In the observation-time PCA, PC1 explained 52.5% of the variance and PC2 explained 30.7% (Fig. S3B). Predictor clustering was overall similar to the treatment-time PCA. Higher values of PC1 correlated with elevation, while lower values correlated with temperature, dew point, vapor point, as well as relative humidity and precipitation. Higher values of PC2 correlated with radiance and higher temperatures, while lower values correlated with precipitation and relative humidity.

*Covariate data and software* **–** Analyses were performed with R version 4.2.2 (R Core Team, 2023) and RStudio version 2023.03.0 (RStudio Team, 2023), using dplyr (Wickham et al., 2021) and sqldf to prepare the data and ggplot2 for visualization (Wickham 2016). Linear mixed effect models were fitted with lme4 (Bates et al., 2015), ordinary kriging was performed using the kriging package (Olmedo, 2022), and PCA was performed using the base R stats library. The car package (Fox et al., 2012) was used to obtain Anova tables for each GLMM. The Adaptive Gauss-Hermite Quadrature was used (with nAGQ=0 in lme4) was used to allow model convergence on both GLMMs in the per-timestep analyses, which can cause less accurate but nonetheless acceptable model fits (Stegmann et al., 2018).

Climate data were directly obtained or derived from DAYMET version 4 (Thornton et al., 2022) using the daymetr package (Hufkens et al., 2018). Standard deviation of elevation was derived from the USGS 3DEP model (10.2m) dataset (U.S. Geological Survey, 2022) and land use was derived from the ESA WorldCover (10m) dataset (Zanaga et al., 2022), using Google Earth Engine version 0.1.329 (Gorelick et al. 2017) via the rgee (Aybar, 2022) and reticulate (Ushey et al., 2023) packages. The Colorado TIGER/Line shapefile was obtained using the tigris package (Walker, 2023). Basemap (Figure S1A) was obtained from Google Maps (Google, 2023).

## 3. Results

### 3.1 Collection and processing of inoculum

During the course of this study, CT patches naturally infected with *Puccinia* were discovered at locations across the state of CO and used as sources of telia-bearing leaves (Fig. 1 C). From 2013-2021, both our scouting and method for inoculum preparation improved, going from an initial ∼1.5 kg of telia-bearing leaves processed in 2013 to ∼18 kg in 2021 (Fig. S2). 2019 was the best year for harvest, yielding ∼33 kg.

### 3.2 Timing of inoculation

There was a 73-day range of optimum inoculation dates, predicted from historic weather data, among sites. The range generally coincided with elevation, and the earliest optimum dates (July 9) were usually at the higher elevations, as opposed to later dates (September 20) at lower elevations. Despite this range, inoculations were broadly successful across the state regardless of exact timing.

### 3.3 Overall stem counts

At most sites (∼77%), CT stem counts declined from initial measurement to final observation, from a mean (± SE) of 87.9 ± 6.5 stems to 44.7 ± 4.2 stems (Figure 1A). Percent change from initial to final stem counts was spatially autocorrelated however (Moran’s I = 0.186 vs. –0.012 expected, p < 0.01, Kriged values presented in Fig. 1B).

### 3.4 Annual Stem Counts

*Treatment and protocol effects* – Stem count at time *t* was influenced by stem count at time *t*-1, monitoring year, last treatment type, last treatment quantity, and years since the last treatment (Table 1). Marginal effects of each predictor are presented in Fig. 3. Stem counts were lower in later monitoring years (Fig. 3A), and higher stem counts at time *t*-1 resulted in lower stem counts at time *t*, suggesting negative density dependence (Fig. 3B). Stem counts were lowest after Broadcast treatment (95% CI: 40.2-59.4), highest after Targeted treatment (95% CI: 52.4-71.0), and intermediate after Targeted+Broadcast (95% CI: 46.2-66.2) (Fig. 3C). Stem counts also increased with increasing time since the last treatment was applied (Fig. 3D), and declined with increasing quantity of inoculum at the last treatment (Fig. 3E).

**Figure 3.**
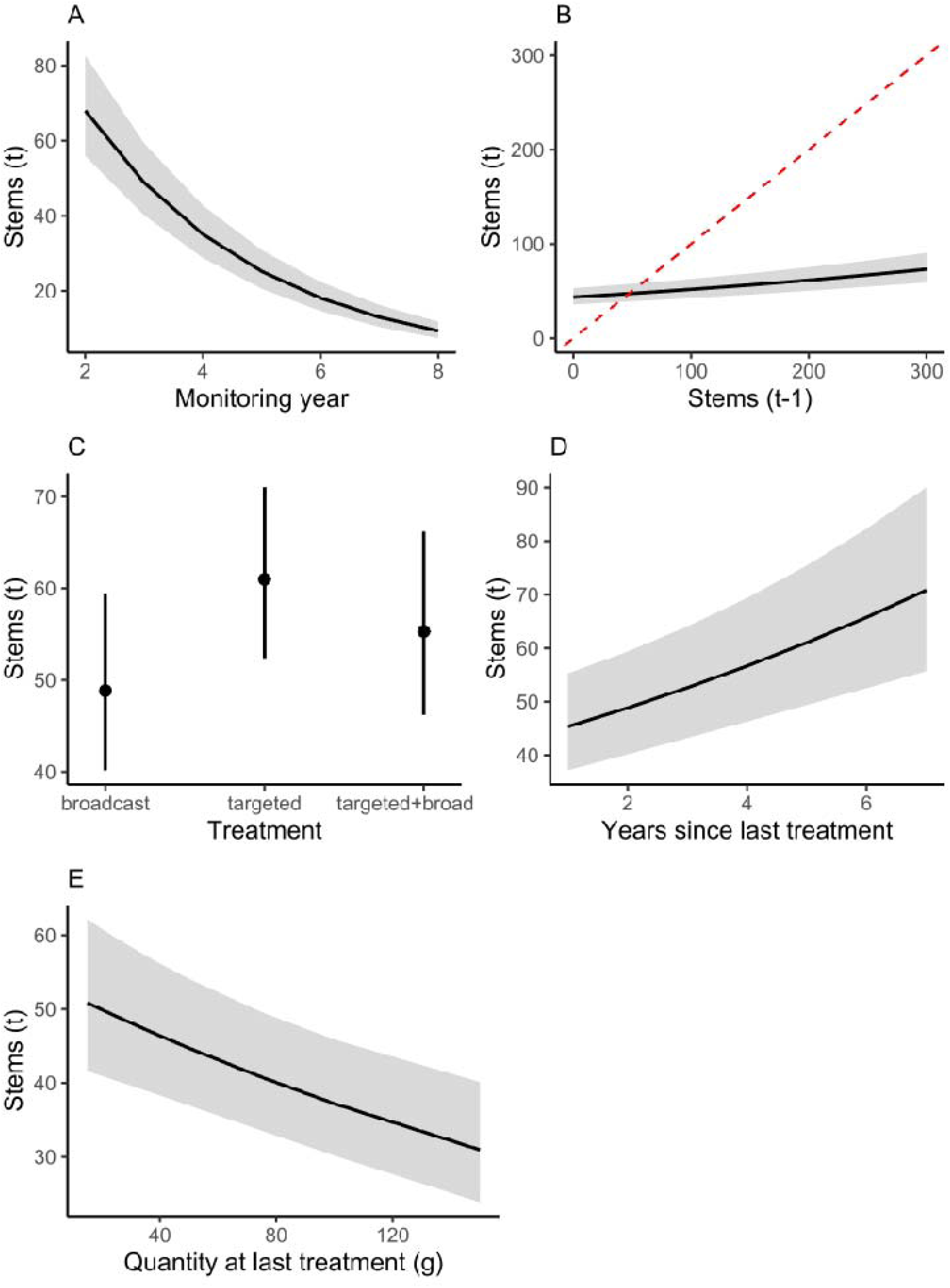
Marginal effects on Canada thistle stem counts at time *t* of (A) monitoring year, (B) stem count in the prior year, (C) last treatment type, (D) years since last treatment, and € quantity of *P. punctiformis* inoculum at last treatment. Shaded areas (and error bars in panel C) represent 95% confidence intervals (*N* = 87). Dashed red line in panel B shows y=x, with values above this line indicating an overall increase in stem counts from the prior year, and values below indicating a decline.

**Table 1.** Poisson GLMM results for Canada thistle stem counts at time *t* as a function of: stem counts at time *t*-1, monitoring year, last treatment type, last treatment quantity (of biocontrol inoculum), and years since last treatment. Site was included as a random intercept term. Statistically significant results (p < 0.05) highlighted in bold.

| Parameter | $\chi^2$ | DF | $p$ |
| --- | --- | --- | --- |
| last_stems | 85.54 | 1 | **< 0.001** |
| monitoring_year | 680.01 | 1 | **< 0.001** |
| yrs_since_trt | 25.44 | 1 | **< 0.001** |
| amount_lasttrt | 16.55 | 1 | **< 0.001** |
| treatment_lasttrt | 12.50 | 2 | **0.002** |

**Table 2.** Poisson GLMM results for Canada thistle stem counts at time *t* as a function of: stem counts at time *t*-1, monitoring year, annual mean maximum temperature, standard deviation of elevation, and PC1 and PC2 from treatment-time PCA and observation-time PCA. Standard deviation of elevation was calculated for a 2.5 radius circle around each site. Site was included as a random intercept term. Statistically significant results (p < 0.05) highlighted in bold.

| Parameter | $\chi^2$ | DF | $p$ |
| --- | --- | --- | --- |
| last_stems | 122.29 | 1 | **< 0.001** |
| monitoring_year | 1325.58 | 1 | **< 0.001** |
| PC1_obs | 27.92 | 1 | **< 0.001** |
| PC1_trt | 18.34 | 1 | **< 0.001** |
| PC2_obs | 4.32 | 1 | **0.038** |
| PC2_trt | 4.64 | 1 | **0.031** |
| tmax_ann | 103.15 | 1 | **< 0.001** |
| elev_stddev_m | 12.80 | 1 | **< 0.001** |
| pct_dist | 0.00 | 1 | 0.975 |

*Climate and geographic effects* – Controlling for stem counts at *t*-1 and monitoring year as covariates, stem counts at time *t* were also affected by mean annual daily maximum temperature and standard deviation of elevation, as well as principal components based on climate data from treatment time and observation time (Table 3). Stem counts at time *t* declined with increasing mean annual daily maximum temperatures (Fig. 4A), and with increasing standard deviation of elevation (i.e., less ‘flatness’, Fig. 4B).

**Figure 4.**
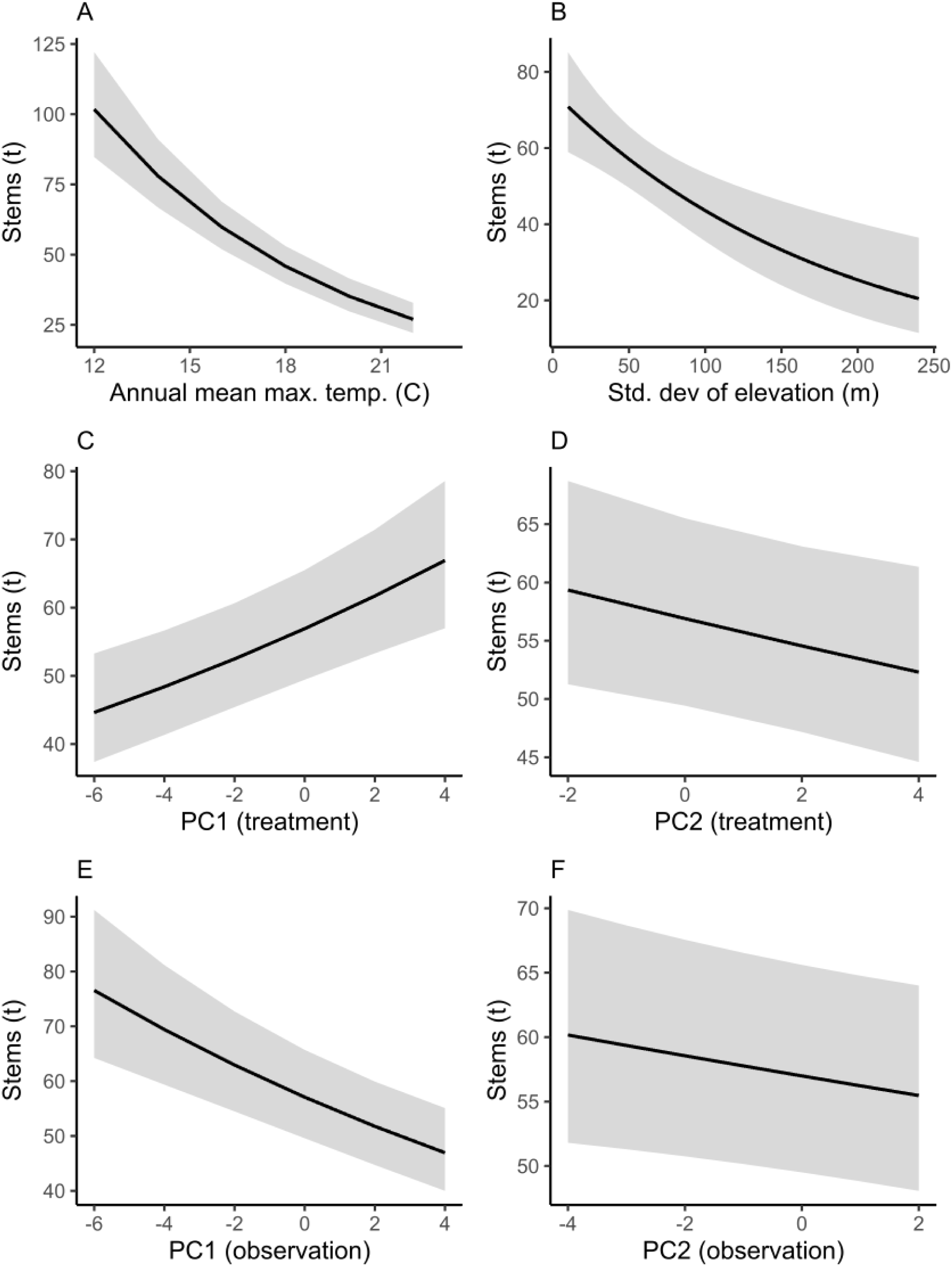
Marginal effects on stem counts at time *t* of (A) annual mean maximum temperature (°C), (B) standard deviation of elevation (2.5 km radius around each site), (C) PC1 from treatment-time PCA, (D) PC2 from treatment-time PCA, (E) PC1 from observation-time PCA, and (F) PC2 from observation-time PCA. Shaded areas represent 95% confidence intervals (*N* = 87).

Stem counts decreased with lower PC1 scores (Fig. 4C) and higher PC2 scores (Fig. 4D) based on the PCA on climate around treatment time. In other words, stem counts were lower where in the 2 weeks prior to treatment sites had: both lower radiance and elevation, and higher temperatures, vapor pressure, dew point, and relative humidity. The effects of precipitation in the 2 weeks prior to treatment were more ambiguous however.

Based on the PCA on climate around observation time, stem counts decreased with higher PC1 (Fig. 4E) and PC2 (Fig. 4F) scores. In other words, stem counts were also lower where, in the 2 weeks prior to monitoring, sites had both a higher radiance and elevation, and lower daily average and minimum temperatures, dew point, vapor point, precipitation, and relative humidity. The effects of daily maximum temperatures however were more ambiguous.

## 4. Discussion

Our efforts suggest that *Puccinia* can be effectively developed as a biological control agent of widespread use for CT management. Collectively, in excess of 125 kg of rust inoculum was harvested and prepared for release, with us successfully treating 87 sites spread across two-thirds of Colorado’s counties. Following these releases, we observed declines in CT stem density in ∼77% of those over the eight-year period. Despite these declines, epiphytotics of systemic rust disease were only observed in ∼17% of the treated sites, verified by the appearance of systemically diseased shoots. Moreover, we observed relationships between the degree of stand decline and the amount of rust inoculum used for treatment, method of treatment application, and time since the last site treatment, suggesting potential for further refinement of our protocol. This measured progress, along with the prospects of *Puccinia* being adopted more widely for use as a biological control agent are discussed in more detail below.

One of our primary objectives in this study was to monitor CT stem densities at inoculation sites and make our findings available to weed managers. We strongly believe that the fungus is the primary causative agent responsible for the CT decline observed. First, the pattern of stand decline detected exhibits all the hallmarks of conspecific negative density dependence (Fig. 2A; CNDD; Chesson 2000), as CT stands with the highest initial density tended to exhibit the most precipitous decline in abundance by the end of our study period. Though other factors can contribute to CNDD (e.g., intraspecific competition), host-specific natural enemies are a primary driver, as has been observed in other tropical and temperate systems (Comita et al. 2012). Second, Canada thistle patches are long-lived and usually expand through extensive root systems, displaying the invasive tendency that is indicated by the common name used in the native range of the plant, creeping thistle (Cockayne, 1915; Moore, 1975; Cheater, 1992; Donald, 1994; Nuzzo, 1997; Morishita, 1999; Guiggisberg, *et al*., 2012). Counter to this invasive tendency, CT stem densities have declined as measured in this and other studies (Berner et al., 2013b; 2015) where patches had been inoculated with *Puccinia*. Furthermore, the rate and extent of decline observed in our study is comparable to those conducted on a much smaller scale in Maryland (U.S.), Greece and Russia (Berner et al., 2015a). Third, we observed significant relationships between the method of treatment employed (Fig. 3C), timing since the last site treatment (Fig. 3D), and the amount of rust inoculum used (Fig. 3E) with the magnitude of CT stem decline. Finally, although there is the possibility that factor(s) other than *Puccinia* may have been involved in contributing to CT decline, as far as we can tell there are no such other factor(s) and we haven’t found accounts of similar regional declines in CT. Recommended conventional procedures for control are difficult and time consuming, involving herbicide applications and mowing over several seasons (Beck and Sebastian, 2000; Beck, 2008) which would make discovery of another factor or factors extremely valuable in CT management, yet none have been found.

Another primary objective of our study was to develop an effective inoculation protocol and reliably produce sufficient inoculum to satisfy a statewide program. Over the course of this study, we developed a network of cooperators, including landowners and weed managers, who have helped us locate systemically diseased CT patches, and allowed us to harvest, process and store telia-bearing leaves in amounts necessary to produce over 20 kg of inoculum annually (sufficient to treat CT at 400 sites, 50 rosettes per). We have also trained landowners to harvest, process and release inoculum collected from their own properties. These efforts have not been limited to CO, as we have engaged agricultural producers and stakeholders in other western states in order to more widely promote this CT biocontrol option. More specifically. *Puccinia* and our inoculation protocol are now being utilized by practitioners in Oregon, Washington, Idaho, Montana, Wyoming, Utah and Nevada, among others. This multi-state program has been facilitated by USDA APHIS 526 permits for movement of CT rust from CO to cooperating partners in these neighboring states (Bean and Randall, 2017); however, movement has ceased for the time being as the regulatory status of *Puccinia* as a biocontrol agent is debated. As a result, we put Colorado’s statewide distribution program on temporary hold in 2021 at a point when demand was high, with over 1,000 requests for the rust from farmers, ranchers and other end users. While we wait for resolution of permitting issues we continue to address questions regarding the biology of the fungus, it’s impact on CT, and how to best use it in an Integrated Pest Management (IPM) program to achieve the greatest possible long-term control in our state and beyond.

Canada thistle density declines occurred in ∼77 % of the treated sites, which could be a result of sporadic inoculation success. The following are some possible reasons. Due to the wide range in optimum inoculation dates in CO, limited personnel and extensive travel, inoculations on optimum dates for some of the sites were probably missed and resulted in failure to establish disease at these sites. In addition, no consideration was given to optimum time of day for inoculation, and sites were inoculated throughout the day depending on constraints imposed by travel schedules and landowner availability. This could be problematic since basidiospores, the spores responsible for establishing systemic root disease (Menzies, 1953), are hyaline and susceptible to being killed by ultraviolet radiation. Inoculation late in the day or early evening might ensure better survival of basidiospores until infection is accomplished.

Trial sites were on private lands where landowners had identified CT as a problem and were seeking ways to control the weed. Landowners had been requested not to interfere with the trial sites, yet we are aware that herbicide applications, mowing and other treatments were utilized at some locations, which is not surprising given the initial desire to control CT and the patience required for a multi-season project. It’s clear that some treatments may have resulted in transient declines in stem densities, thus confounding observable effects of the rust. Although it seems intuitive that treatments such as herbicide application would cause a decline in CT stem densities, more subtle effects of herbicides could include an inhibition of rust fungus establishment and impact. For instance, systemic herbicides may have substantial fungicidal activity (Berner et al., 1992), and so treatment could possibly kill the fungus at an early stage following inoculation. Clearly, recommendations for use of the rust fungus in conjunction with chemical, mechanical and cultural strategies are sorely needed to understand potential synergies and limitations of combining approaches in an IPM program for CT control (Chichinski et al., 2023).

Establishment of systemic disease can most likely be improved by inoculating at projected optimum dates, inoculating in the evening, and avoiding systemic herbicides. In addition, development of more intensive inoculation protocols could be useful. These might include more frequent releases at a site during a given fall, and over consecutive seasons, which may be necessary given patterns of CT decline observed in relation to the timing since a site was last treated. Limited availability of *Puccinia* teliospores has long been recognized as a factor hampering the use of the fungus in CT control (Cockayne, 1915) but we have been able to collect and process kilograms of inoculum and are training landowners and weed managers to do the same. Limited teliospore availability shouldn’t then hamper development of more intensive inoculation protocols.

A more detailed characterization of the inoculation material and the interaction of infective spores with CT plants could be used to increase the likelihood of establishment, and possibly virulence of the fungus. This would include routine measurements of germination rates of teliospores (Clark et al., 2020) and characterization of the fungus at a molecular level, where it appears that multiple strains exist with the potential for variable impacts on Canada thistle (Bradshaw et al., 2023). It has also been shown that treatment with plant growth regulators may increase the likelihood of infection of Canada thistle by the rust fungus (Clark et al., 2020) and such treatments could be incorporated into inoculation protocols.

A better understanding of the dynamics and impact of *Puccinia* on underground CT structures would also be useful for biocontrol practitioners who need to explain to landowners why a treatment may or may not work for them. It is clear that at some sites (Fig. S4) the fungus spreads rapidly and takes down CT patches completely, or almost so. Olive (1913) stated that the main method of distribution of the disease occurred through creeping rootstocks and interconnectedness of diseased plants through their rootstocks. With this in mind, it is useful to consider *Puccinia* as a parasitic fungus with an extensive mycelium hosted within the root systems of CT. From this perspective, one of the major challenges in evaluating the impact of *Puccinia* is determination of the belowground status of the fungus. In a previous study, an immunochemical detection method was used to show that the fungus was present as an asymptomatic infection in the roots of 50-60% of teliospore-inoculated CT rosettes (Berner et al., 2015a), reinforcing the idea that root decline, the most agriculturally important impact of the fungus, may take place in the absence of the visible, aboveground signs of the pathogen. While useful, the immunochemical method would be difficult to deploy for widespread and high-throughput detection of root infections. A method based on amplification of *Puccinia-*specific DNA using either polymerase chain reaction (PCR) or loop-mediated isothermal amplification (LAMP) would be more useful, but development has thus far been limited (but see Berner et al., 2015a, Henderson et al., 2019; Clark et al., 2020). The potential benefits of a rapid, molecular-based detection method would be well worth the modest research investment needed to develop, and the initial success of our program should provide impetus for further work to measure the establishment and associated impact of the rust belowground.

The ability to recognize fungal presence would also be enhanced by considering infection symptoms beyond the appearance of aboveground spore-bearing structures. Notably, other disease symptoms include a decrease in stem height where the fungus is present below ground (Berner et al., 2015a; Fig S5) as well as flower bud desiccation and red stems noted during this study, and seen previously (Berner et al., 2015a). These should be characterized in additional detail, and their measurement incorporated into monitoring protocols. It is almost certain that CT plants systemically diseased with *Puccinia* would have a spectral signature associated with the disease, including the stress of a declining root system, which could be detected by one or more of several imaging options used for detection of plant disease (Mahlein, 2016; Lowe et al., 2017). Implementation of detection methods based on optical or other imagery would inform landowners of *Puccinia* presence and extent, informing decisions regarding CT management. Given the high likelihood of success and the economic importance of CT in N. America, development of imaging techniques to detect presence and extent of patch infection would be valuable.

The disease associated with *Puccinia* infection of CT roots has not, to our knowledge, been named. The fungus is commonly referred to as *C. arvense* rust or CT rust. These names are not descriptive of the disease, which is primarily manifested in root decline and host death, and can be misleading to end users in particular who may focus on the foliar aeciospore (rust) stage, which may not occur every year, and fail to note the more agriculturally important decline in thistle stem density and decreasing thistle vigor. In addition to shoot density decline, CT plants may display symptoms consistent with declining root function. Older symptomatic shoots are short, frequently with red stems, with dying flowers and capitula and immunochemical measurements confirmed the presence of the fungus in roots of shoots with these symptoms, even without the appearance of systemically diseased shoots (Berner et al., 2015a). Dieback of shoots and development of stem-free areas within thistle patches follow initial symptoms (Fig. S5), and thistle shoot density decline within diseased patches is a concomitant symptom. One or more of these symptoms have been consistently observed in thistle patches diseased by the fungus. Over several years, dieback spreads to all of the roots and emerging shoots in the patches and the diseased patches ultimately die (Berner et al., 2015a). We propose the name of *Cirsium arvense* dieback (CADB) for the disease and suggest that disease progress be evaluated based on the extent of the aforementioned symptoms of root mortality.

## 5. Conclusions

A statewide CT biocontrol program was successfully implemented in Colorado, U.S.A, using the naturalized rust fungus *Puccinia* as a biocontrol agent. Field scouting, combined with the participation of farmers, ranchers and weed control professionals, enabled the CDA to collect and process sufficient amounts of infected CT leaves to produce more than 20 kg of inoculum annually, enough to inoculate ∼400 sites using the current protocol. Since the disease caused by *Puccinia* has not been named, we propose that it be called *Cirsium arvense* dieback (CADB). This name is useful in that it is based on the visible and agriculturally important disease symptoms (compromised health of the aboveground plant and eventual decline of stem densities), which are an indication of the expansion of the fungus within the root systems of diseased plants, eventually leading to plant death.

## Availability of data and materials

Data and associated scripts are publicly available on Github: https://github.com/ecodata-technology/thistlebiocontrol2023.

## Acknowledgments

We would like to acknowledge the contributions of biocontrol technicians and seasonal staff members to this project, particularly in monitoring field sites across Colorado and collecting teliospore-bearing plant material. These staff members include Sonya Daly, Michael Racette, John Gholson, and Amanda Stahlke. We would also like to acknowledge the cooperation and participation of Colorado’s landowners, who have graciously allowed us access to their properties during these studies.

## References

1. Amor, R. L., and R. V. Harris. 1975. Seedling establishment and vegetative spread of *Cirsium arvense* (L.) Scop. in Victoria, Australia. Weed Research 15:407–411.

2. Amor, R.L. and R.V. Harris. 1977. Control of Canada thistle *Cirsium arvense* (L) Scop. by herbicides and mowing. Weed Research. 17: 303–309.

3. Ayber, C. (2022) rgee: R bindings for calling the ‘Earth Engine’ API. R package version 1.1.5, https://CRAN.R-project.org/package=rgee

4. Bates, D., Mächler, M., Bolker, B. & Walker, S. (2015). Fitting Linear Mixed-Effects Models Using lme4. Journal of Statistical Software, 67, 1–48.

5. Bean, D. and C. Randall. 2017. Colorado and the US Forest Service initiate a multistate Canada thistle biocontrol project. PP. 44. Western Society of Weed Science Newsletter, Fall 2017.

6. Beck, G. K. 2008. Canada thistle. Colorado State University Cooperative Extension, Fact Sheet No. 3.108: 1-3.

7. Beck, K.G. and J.R. Sebastian. 2000. Combining mowing and fall-applied herbicides to control Canada thistle (*Cirsium arvense*). Weed Technology, 14: 351–356.

8. Berner, D. K., E. L. Smallwood, C. A. Cavin, M. B. McMahon, K. M. Thomas, D. G. Luster, A. L. Lagopodi, J. N. Kashefi, Zhanna Mukhina, Tamara Kolomiets, Lyubov Pankratova. 2015a. Asymptomatic systemic disease of Canada thistle (*Cirsium arvense*) caused by *Puccinia punctiformis* and changes in shoot density following inoculation. Biological Control 86: 28–35.

9. Berner, D. K., Smallwood, E. L., Vanrenterghem, M., Cavin, C. A., Michael, J. L., Shelley, B. A., … & Mukhina, Z. 2015b. Some dynamics of spread and infection by aeciospores of *Puccinia punctiformis*, a biological control pathogen of *Cirsium arvense*. Biological Control, 88, 18–25.

10. Berner, D. K. and D.W. Bean. 2013a. Biological control of Canada thistle using a rust fungus and releasing APHIS-approved pathogens for control of Russian knapweed and Russian thistle. USDA, ARS and Colorado Dept. of Ag. Specific Cooperative Agreement, ARS Project Number 8044-22000-039-07

11. Berner, D. K., E. Smallwood, C. Cavin, A. Lagopodi, J. Kashefi, T. Kolomiets, L. Pankratova, Z. Mukhina, M. Cripps, G. Bourdôt. 2013b. Successful establishment of epiphytotics of *Puccinia punctiformis* for biological control of *Cirsium arvense*. Biological Control 67:350–360.

12. Berner, D. K., G. T. Berggren, and J. P. Snow. 1992. Method for protecting plants against soil-borne fungi using glyphosate and imazaquin compositions. United States Patent Number 5,110,805)

13. Bradshaw, M.J., Carey, J., Liu, M., Bartholomew, H.P., Jurick, W.M., Hambleton, S., Hendricks, D., Schnittler, M., M. Scholler (2023) Genetic time traveling: sequencing old herbarium specimens, including the oldest herbarium specimen sequenced from kingdom Fungi, reveals the population structure of an agriculturally significant rust. New Phytologist 237: 1463–1473.

14. Buller, A.H.R.1950. Researches on fungi. Vol. VII: The sexual process in Uredinales. The University of Toronto Press: Toronto. Pp 344–388.

15. Carlson, S. J. and W. W. Donald. 1988. Fall-applied glyphosate for Canada thistle (*Cirsium arvense*) control in spring wheat (*Triticum aestivum*). Weed Tech. 2: 445–455.

16. Cheater, M. 1992. Alien invasion. Nature Conservancy 42:24–29.

17. Chichinsky D, Larson C, Eberly J, Menalled FD and Seipel T (2023) Impact of *Puccinia punctiformis* on *Cirsium arvense* performance in a simulated crop sequence. Front. Agron. 5:1201600. doi: 10.3389/fagro.2023.1201600

18. Clark, A. L., Jahn, C. E., and Norton, A. P. (2020). Initiating plant herbivory response increases impact of fungal pathogens on a clonal thistle. Biol. Control 143, 104207. doi: 10.1016/j.biocontrol.2020.104207

19. Cockayne, A.H. 1915. Californian thistle rust. J. Agric. 12: 300–302.

20. Connick, W.J., and R.C. French. 1991. Volatiles emitted during the sexual stage of the Canada thistle rust fungus and by thistle flowers. J. Agric. Food Chem. 39:185–188.

21. Cripps, M.G., A. Gassmann, S.V. Fowler, G.W. Bourdot, A.S. McClay, and G.R. Edwards (2011) Classical biological control of Cirsium arvense: Lesions from the past. Biological Control 57: 165–174

22. Cripps, M. G., G. W. Bourdot, K. L. Bailey. 2012. Plant pathogens as biocontrol agents for *Cirsium arvense* – an answer to Muller and Nentwig. Neobiota 13: 31–39.

23. Donald, W.W. 1994. The biology of Canada thistle (Cirsium arvense) Reviews of Weed Science 6:77–101.

24. Ferdinandsen, C. 1923. Biologiske Undersogelser over Tidsel rust (*Puccinia suaveolens* (Pers.) Rostr.). Nordisk Joudbrugs-forskning. 5-8:475-487.

25. Fox, J., Weisberg, S., Adler, D., Bates, D., Baud-Bovy, G., Ellison, S., … & Heiberger, R. 2012. Package ‘car’. Vienna: R Foundation for Statistical Computing, 16.

26. French, R. C. (1990). Stimulation of germination of teliospores of *Puccinia punctiformis* by nonyl, decyl, and dodecyl isothiocyanates and related volatile compounds. J. Agric. Food Chem. 38, 1604–1607. doi: 10.1021/jf00097a037

27. French, R.C. and A.R. Lightfield. 1990. Induction of systemic aecial infection in Canada thistle (*Cirsium arvense*) by teliospores of *Puccinia punctiformis*. Phytopathology. 80:872–877.

28. Google. (n.d.) [Google Map of Colorado with sites pinned]. Retrieved June 20, 2023.

29. Gorelick, N., Hancher, M., Dixon, M., Ilyushchenko, S., Thau, D., & Moore, R. (2017). Google Earth Engine: Planetary-scale geospatial analysis for everyone. Remote Sensing of Environment, 202, 18–27.

30. Guiggisberg, A., E. Welk, R. Sforza, D. P. Horvath, J. V. Anderson, M. E. Foley, L. M. Rieseberg,. 2012. Invasion history of North American Canada thistle, *Cirsium arvense*. Journal of Biogeography 39: 1919–1931.

31. Halsted, B. D. 1894. Weeds and their most common fungi. New Jersey Expt. Sta. Rept. 379–381.

32. Hamdoun, A.M. 1970. The anatomy of subterranean structures of Cirsium arvense (L.) Scop. Weed Research. 10:284–287.

33. Henderson, C., Cripps, M., & Casonato, S. (2019). Distribution of *Puccinia punctiformis* in above-ground tissue of *Cirsium arvense* (Californian thistle). New Zealand Plant Protection, 72, 265–270.

34. Hufkins, K., Basler, D., Milliman, T., Melaas, E.K., & Richardson, A.D. (2018) An integrated phenology modelling framework in R: modelling vegetation phenology with phenor. Methods in Ecology & Evolution, 9, 1–10.

35. Jacobs, J., J. Sciegienka, and F. Menalled. 2006. Ecology and Management of Canada thistle *Cirsium arvense* (L.) Scop. United States Department of Agriculture NATURAL Resources Conservation Service Invasive Species Technical Note No. MT-5.

36. Kartesz, J.T. 2015. The Biota of North America Program (BONAP). 2015. Taxonomic Data Center. (http://www.bonap.net/tdc). Chapel Hill, N.C.

37. Kay, Q.O.N. 1985. Hermaphrodites and subhermaphrodites in a reputedly dioecious plant, *Cirsium arvense* (L.) Scop. New Phytologist. 100:457–472.

38. Lowe, A., N. Harrison and A.P. French (2017) Hyperspectral image analysis techniques for the detection and classification of the early onset of plant disease and stress. Plant Methods 13:80 DOI 10.1186/s13007-017-0233-z

39. Magnusson, M.U., D.L. Wyse, and J.M. Spitzmueller. 1987. Canada thistle (*Cirsium arvense*) propagation from stem sections. Weed Science. 35:637–639.

40. Mahlein, A-K (2016) Plant disease detection by imaging sensors – parallels and specific demands for precision agriculture and plant phenotyping. Plant Disease DOI 10.1094/PDIS-03-15-0340-FE

41. Menzies, B.P. 1953. Studies on the systemic fungus, *Puccinia suaveolens*. Annals of Botany. 17:551–568.

42. Moore, R. J. 1975. The biology of Canadian weeds. Canadian Journal of Plant Science 55:1033–1048.

43. Morishita, D.W. 1999. Canada thistle. pages 162-172 In: Sheley and Petroff (ed.), Biology and management of noxious rangeland weeds. Oregon State University Press.

44. Naef, A., Roy, B.A., Kaiser, R., Honegger, R., 2002. Insect-mediated reproduction of systemic infections by *Puccinia arrhenatheri* on *Berberis vulgaris*. New Phytol.

45. Nuzzo, V. 1997. Element stewardship abstract for Cirsium arvense. The Nature Conservancy 1–30.

46. Olive, E. W. 1913. Intermingling of perennial sporophytic and gametophytic generations in *Puccinia podophylii*, P. obtegens and Uromyces glycyrrhizae. Annales Mycologici 11: 297–311.

47. Olmedo, O.E. (2022). kriging: Ordinary Kriging. R package version 1.2, https://CRAN.R-project.org/package=kriging

48. Price, J.R. 2014. Post release assessment of classical biological control of Canada thistle (Cirsium arvense) in the western United States. MS Thesis, 130 pp., Department of Plant, Soil and Entomological Sciences, University of Idaho, Moscow, ID.

49. Rostrup, F. G. E. 1874. Ein eigenthumliches generations-verhaltniss bei *Puccinia sauveolens* (Pers.). Bot. Ztg. 32: 556–557.

50. R Core Team. (2023). R: A Language and Environment for Statistical Computing. R Foundation for Statistical Computing, Vienna, Austria.

51. RStudio Team. (2023). RStudio: Integrated Development Environment for R. RStudio, PBC, Boston, MA.

52. Stachon, W. J., and R. L. Zimdahl. 1980. Allelopathic activity of Canada thistle (*Cirsium arvense*) in Colorado. Weed Science 28:83–86.

53. Stegmann, G., Jacobucci, R., Harring, J.R., & Grimm, K.J. (2018). Nonlinear mixed-effects modeling programs in R. Structural Equation Modeling, 25, 160–165. Available at: 10.1080/10705511.2017.1396187.

54. Thomas, R.F., T.J. Tworkoski, R.C. French and G.R. Leather. 1994. *Puccinia punctiformis* affects growth and reproduction of Canada thistle (*Cirsium arvense*). Weed Technol. 8, 488–493.

55. Thornton, M.M., R. Shrestha, Y. Wei, P.E. Thornton, S-C. Kao, & B.E. Wilson. (2022). Daymet: Daily Surface Weather Data on a 1-km Grid for North America, Version 4 R1. ORNLDAAC, Oak Ridge, Tennessee, USA. 10.3334/ORNLDAAC/2129

56. Turner, S. K., P. K. Fay, E. L. Sharp, and D. Sands. 1981. Resistance of Canada thistle (*Cirsium arvense*) ecotypes to a rust pathogen (*Puccinia obtegens*). Weed Science 29:623–624.

57. U.S. Geological Survey. (2022). 3D Elevation Program 10-Meter Resolution Digital Elevation Model. https://www.usgs.gov/3d-elevation-program

58. Ushey, K., Allaire, J., Tang, Y. (2023). reticulate: Interface to ‘Python’. R package version 1.29, https://CRAN.R-project.org/package=reticulate

59. Van den Ende, Q.J., J. Frantzen, and T. Timmers. 1987. Teleutospores as origin of systemic infection of *Cirsium arvense* by *Puccinia punctiformis*. Netherlands Journal of Plant Pathology. 93:233–239.

60. Walker, K. (2023). tigris: Load census TIGER/Line shapefiles. R package version 2.0.3, https://CRAN.R-project.org/package=tigris

61. Watson, A.K. and W.J. Keogh. 1980. Mortality of Canada thistle due to Puccinia punctiformis. Proc. V Int. Symp. Biol. Contr. Weeds, Brisbane, Australia: 325–332.

62. Wickham, H. (2016). ggplot2: Elegant Graphics for Data Analysis. Springer-Verlag New York.

63. Wickham, H., François, R., Henry, L. & Müller, K. (2021). dplyr: A Grammar of Data Manipulation. R package version 1.1.0, https://CRAN.R-project.org/package=dplyr

64. Wilson, C. L. 1969. Use of plant pathogens in weed control. Pages 411–434. In: Horsfall, J. (ed.), Ann. Rev. Phytopathology **7:** Annual Reviews, Inc., Palo Alto.

65. Winston, R. L., Schwarzländer, M., Hinz, H. L., Day, M. D., Cock, M. J., & Julien, M. H. (2014). Biological control of weeds: A world catalogue of agents and their target weeds. Biological control of weeds: a world catalogue of agents and their target weeds., (Ed. 5).

66. Zanaga, D., Van De Kerchove, R., Daems, D., De Keersmaecker, W., Brockmann, C., Kirches, G., Wevers, J., Cartus, O., Santoro, M., Fritz, S., Lesiv, M., Herold, M., Tsendbazar, N.E., Xu, P., Ramoino, F., & Arino, O. (2022). ESA WorldCover 10 m 2021 v200. doi:10.5281/zenodo.7254221

