## Supplementary Information for "Scaling use of the rust fungus *Puccinia punctiformis* for biological control of Canada thistle (*Cirsium arvense* (L.) Scop.): First report on a U.S. statewide effort"

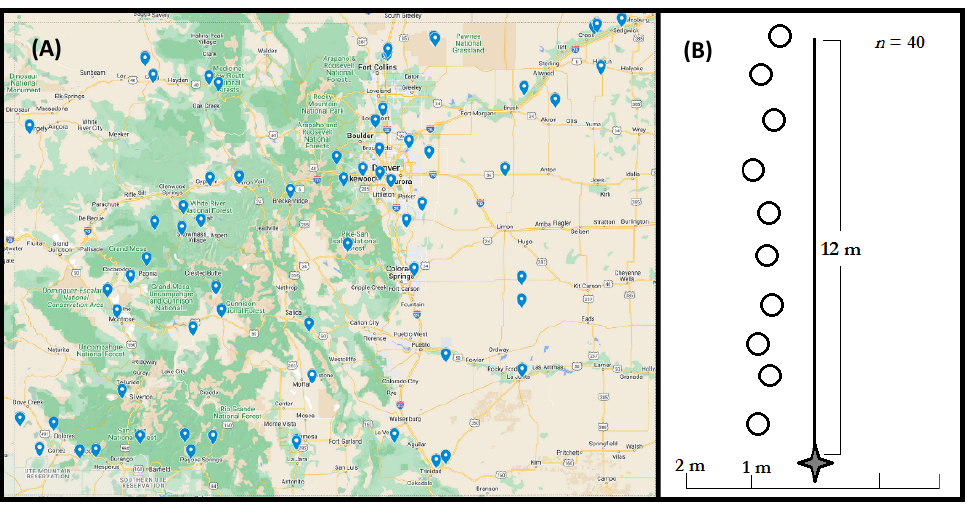

**Figure S1.** Map (A) of Colorado sites (*N* = 87; blue pins) invaded by Canada thistle and treated with *Puccinia punctiformis* teliospores. At each site, a permanent, 12 x 4 m transect (B) was established for both inoculating Canada thistle and post-treatment monitoring of disease development and stand dynamics.

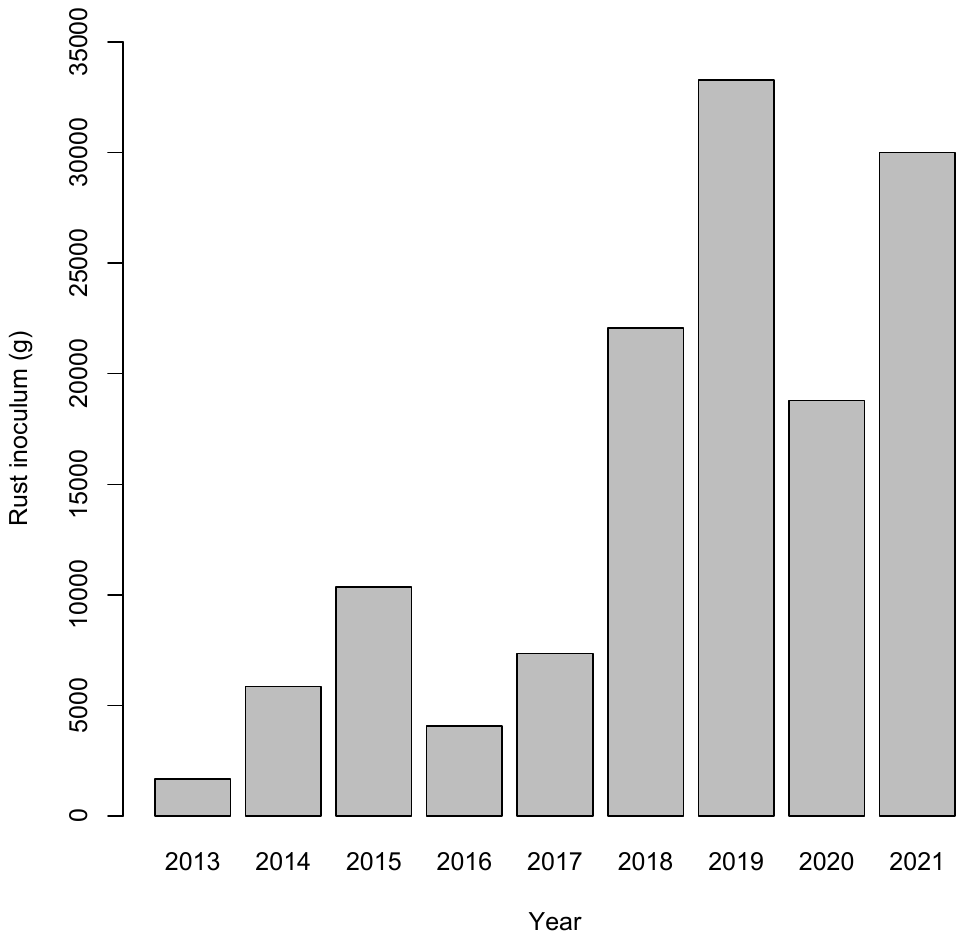

**Figure S2.** Amount (g) of *Puccinia punctiformis* inoculum harvested and processed by the Colorado Department of Agriculture from 2013-2021. Teliospores were either released in the same year, or frozen and used in the subsequent season.

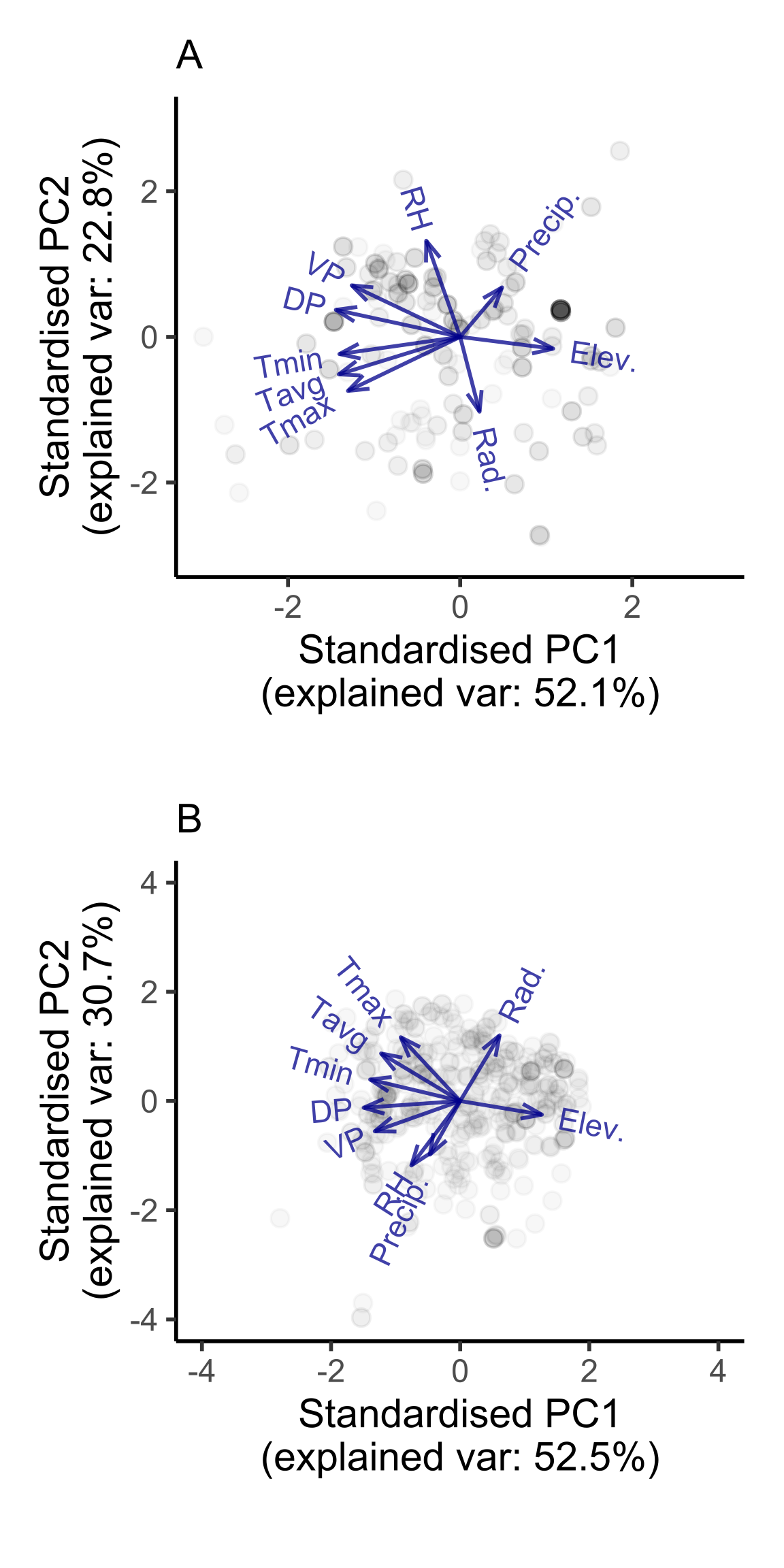

**Figure S3.** Biplots of A. treatment-time PCA, and B. observation-time PCA. Points represent site-year combinations used to train the PCAs, arrows represent loadings for each predictor. Tmax = daily maximum temperature, Tavg = daily average temperature, Tmin = daily minimum temperature, Rad = radiance, VP = vapor pressure, RH = relative humidity, DP = dew point, Precip = precipitation, Elev = elevation.

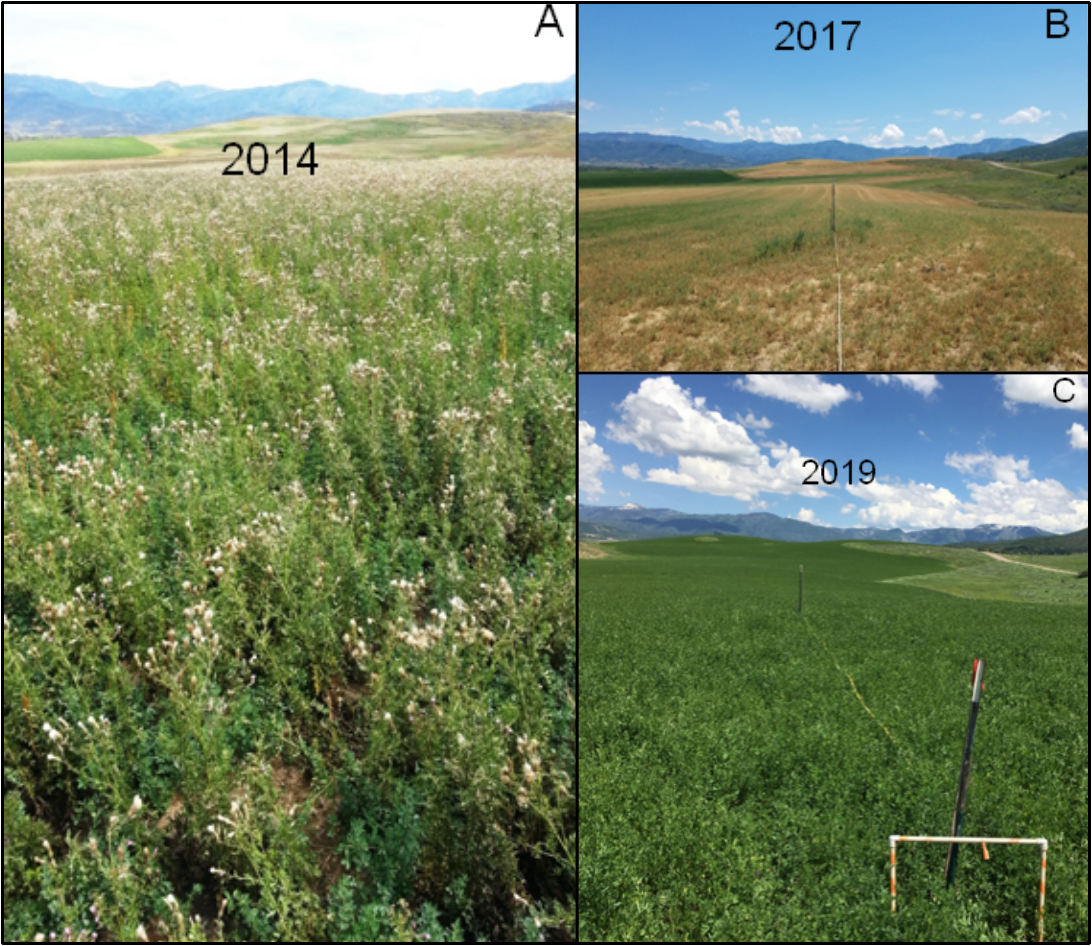

**Figure S4.** Clay site in Colorado showing the progression of Canada thistle decline across a portion of our study period: (A) heavy infestation before rust inoculation, (B) thirty months post-inoculation, there was a 99% reduction in thistle stem density, and (C) same site 54 months after inoculation, with an alfalfa crop and no Canada thistle. Photo credits: A. Joel Price, B&C. Karen Rosen

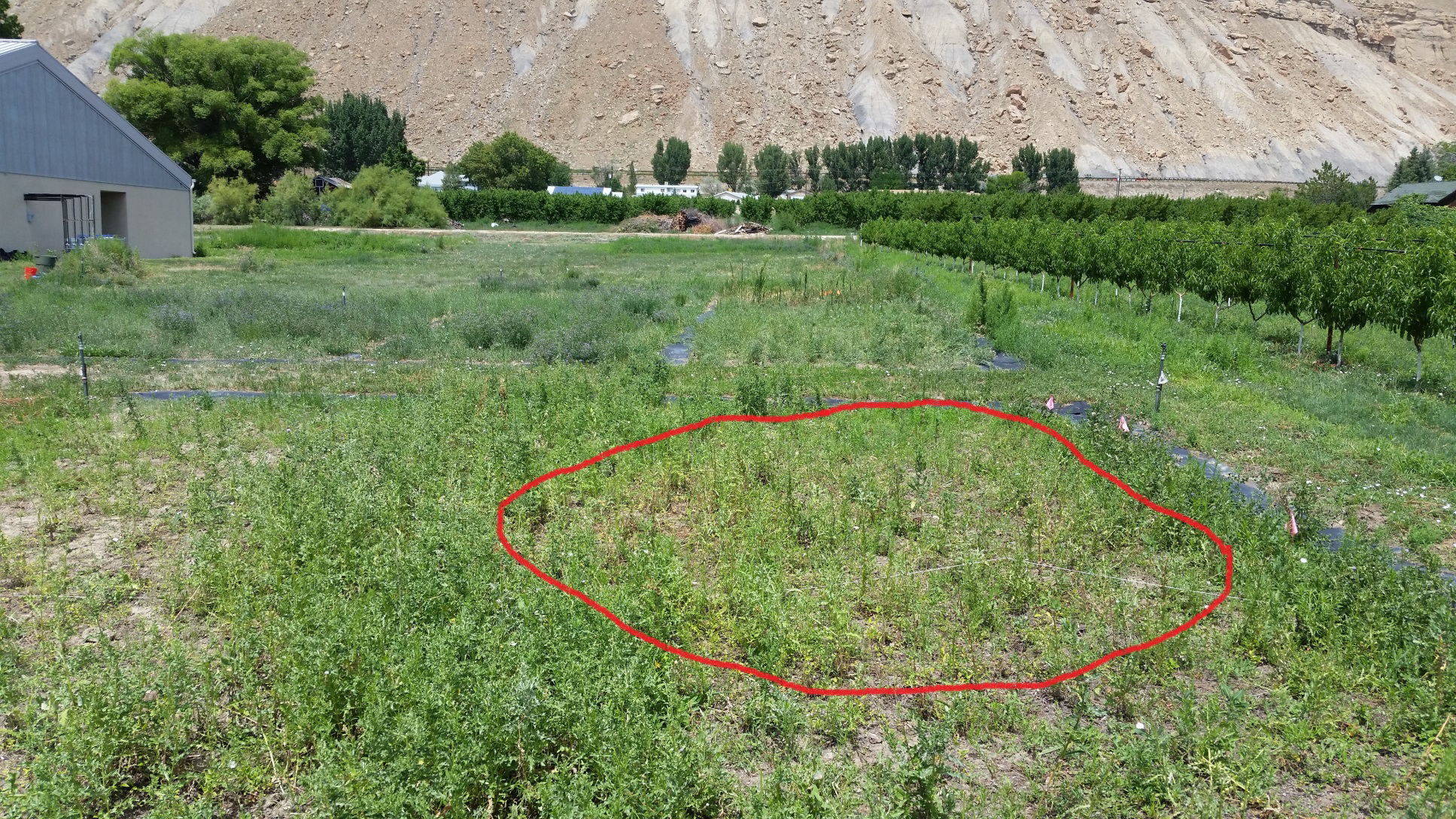

**Figure S5.** Dieback of Canada thistle shoots and development of stem-free areas within an inoculated thistle patch at the Colorado Department of Agriculture experimental garden, indicators of systemic root disease caused by *Puccinia punctiformis.* Photo credit: Joel Price
